# Dnmts mediate neural processing after odor learning in the honeybee

**DOI:** 10.1101/056333

**Authors:** Stephanie D. Biergans, Charles Claudianos, Judith Reinhard, C. Giovanni Galizia

**Affiliations:** Queensland Brain Institute, The University of Queensland, Australia; Neurobiologie, Universitat Konstanz, Germany; Monash Institute of Cognitive and Clinical Neuroscience, Faculty of Medicine, Nursing Health and Sciences, Monash University, Australia

## Abstract

DNA methyltransferases (Dnmts)-epigenetic writers catalyzing the transfer of methyl-groups to cytosine-regulate stimulus-specific olfactory long-term memory (LTM) formation and extinction in honeybees. The physiological relevance of their function in neural networks, however, remains unknown. Here, we investigated how Dnmts impact neuroplasticity in the bees’ primary olfactory center, the antennal lobe (AL) an equivalent of the olfactory bulb of vertebrates. The AL is crucial for odor discrimination, an indispensable process in stimulus-specific LTM. Using pharmacological inhibition, we show that Dnmts promote fast odor pattern separation in trained bees. We show Dnmt activity during memory formation increases both the number of responding glomeruli and the response magnitude to a novel odor. These data suggest that Dnmts are necessary for a form of homoeostatic network control which might involve inhibitory interneurons in the AL network and demonstrate that Dnmts influence neural network properties during memory formation *in vivo*.

## Introduction

The morphology and physiology of the neural network underlying olfactory processing and memory formation has been studied in great detail in honey bees^1^. In the primary olfactory center (antennal lobe, AL), odor information is coded in a spatiotemporal pattern of glomerular activity, which suggests a crucial role of the AL in odor identity processing. Indeed, the representations of individual odors are more distinct after processing in the AL^2^. AL processing is accomplished primarily by the network of inhibitory local interneurons (LNs), as shown by modelling^3^ and by using GABA receptor blockers^4–6^. Odor response patterns separate fast and reach their maximum discriminability about 150 ms after odor onset in the AL output neurons (projection neurons, PNs)^7,8^. Behavioral and physiological studies suggest that bees indeed use this early information for odor discrimination^9–12^. The AL is also involved in olfactory memory formation^13–17^.

Even though olfactory memory formation in bees has been extensively studied at both the physiological and behavioral level, many molecular aspects are poorly understood. Particularly, the dynamics of transcriptional regulation that impact neural processing and underpin memory formation remain largely unknown. Recent studies have shown DNA methylation catalyzed by DNA methyltransferases (Dnmts) regulate stimulus-specific long-term memory (LTM) formation^18,19^ and extinction^20^. *Dnmts* and Ten-eleven translocation methylcytosine dioxygenase *(Tet),* which catalyzes active demethylation, were found upregulated in a specific temporal order following olfactory reward conditioning^19^, highlighting a dynamic relationship between methylation and demethylation. We proposed earlier that Dnmts may normalize transcription levels of genes activated during memory formation, in order to avoid excess neural activity and connectivity^19^. This hypothesis proposes, in essence, homoeostatic plasticity, a slow cell- and/or neural network-wide form of neural plasticity. Similarly, Tet-mediated active demethylation is involved in synaptic scaling, a mechanism of homeostatic plasticity^21,22^.

To better understand how DNA methylation mediates learning-related plasticity in neural networks, we investigated odor responses in the AL output neurons (PNs) with and without Dnmt activity during memory formation, *in vivo.* Dnmt inhibition with RG108 impaired odor response pattern separation between a trained and a new odor. Furthermore, the overall number of glomeruli responsive to a new odor was reduced after Dnmt inhibition, as well as their response strength. Interestingly, inhibiting Dnmts did not change the response to the learned odor. These results suggest an involvement of Dnmts in regulating plasticity in the inhibitory neural network of the AL during memory formation; specifically, the inhibitory network may use Dnmts to undergo compensatory homoeostatic plasticity after olfactory reward conditioning.

## Results

### Dnmts promote stimulus-specific memory formation in bees

Behavioral studies in bees have shown that Dnmts are involved in stimulus-specific LTM^18,19^. When Dnmts were active following olfactory reward conditioning, stimulus-specific memory increased and bees generalized less to a novel odor. But which neural network properties are regulated by Dnmts during LTM formation? In this study, we focused on the primary olfactory center (antennal lobe, AL) of bees. We hypothesized that Dnmts mediate learning-related plasticity in the AL and thus strengthen stimulus-specific memory formation in this neuropil. We combined the use of a Dnmt inhibitor (RG108) with *in vivo* Ca^2+^-imaging of the AL output neurons (projection neurons, PNs). Bees were treated with the inhibitor or the solvent control (DMF) 2 hours after olfactory reward conditioning (Fig. 1a). We tested two behavioral groups, paired and unpaired. In paired training there was a 2s overlap between conditioned (CS) and unconditioned stimulus (US), and a training trial every 10 minutes (inter-trial interval, ITI). In unpaired training, there was a 5 minute gap between CS and US, with the same number, types and rhythm of stimuli as in the paired group, thus serving as stimuli control. We assessed the effect of Dnmt inhibition in paired and unpaired groups for associative or non-associative memory. Bees were trained, stained with the calcium sensitive dye FURA on day 2, and tested on day 3 (Fig. 1a).

**Figure 1:**
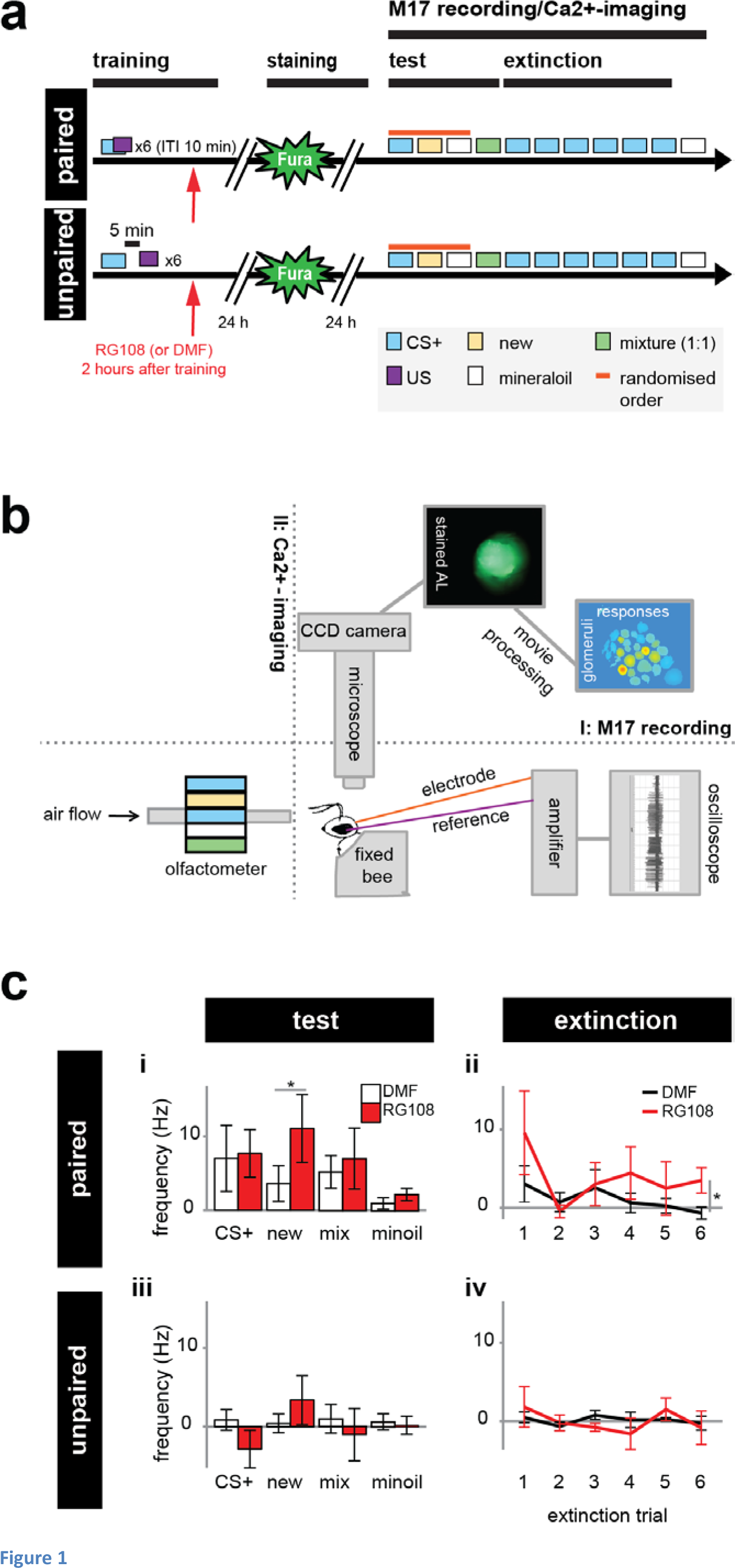
RG108 treatment impairs stimulus-specific memory and extinction in bees. (a) 2 hours after either paired or unpaired training bees were treated with the Dnmt inhibitor RG108 or the solvent DMF (red arrow). 1 day after the training PNs were stained with a Ca^2^+-sensitive dye (Fura). 1 day following the staining, bees were exposed to odors (b) while either their M17 or AL activity was recorded. First, bees were exposed to the trained odor (CS+), a new odor and the odor solvent mineraloil (min) in randomized order followed by the binary mixture of CS+ and new. After that bees received extinction training (6x CS+). (b) Bees’ AL was imaged using a fluorescence microscope with an attached CCD camera. In a separate group of bees the PER muscle (M17) activity was recorded as a behavioral control of Dnmt inhibition efficiency. (c) The M17 spike frequency (mean +/− SEM) is shown 2 days after conditioning. RG108 treated bees responded significantly more to the new odor compared to solvent treated bees (glm interaction treatment, group, stimulus: p=0.038, effect size (d)=0.464). Extinction learning was impaired over time in the paired group (glm interaction treatment, group: p=0.019, d=0.243). Number of bees: paired n(RG108)=19, n(DMF)=26; unpaired n(RG108)=15, n(DMF)=13;* is p<0.05

We recorded learning scores with electrophysiological recordings from the bees proboscis muscle (M17) - in a separate experiment-2 days after conditioning in order to confirm the behavioral effect of Dnmt inhibition (Fig. 1b). We use M17 activity as a proxy for proboscis extension, as done before^23^. In the test phase, we found that control bees responded to the trained odor with a strong M17 response (mean firing rate during 4s stimulus: 7Hz ±4.5Hz), but with no response to the empty stimulus mineral oil (1Hz, ±0.8Hz), showing that the bees had learned to respond to the trained odor (Fig. 1c: i, generalized linear model (glm), interaction treatment, stimulus, group: p = 0.035, effect size (d)=0.435). There was no effect of the Dnmt-inhibitor RG108 on the CS+ response (Fig. 1c: i, glm interaction treatment, stimulus, group: p=0.849), confirming previously published results^18^. As expected, there was no learning in the unpaired group for either treatment (Fig. 1c: iii, glm interaction treatment, stimulus, group: DMF: p=0.957, RG108: p=0.382). Additionally, we tested the bees' responses to a new odor in order to test for stimulus-specific memory and generalization. We found that RG108-treated bees generalized significantly more to a new odor compared to control bees (Fig. 1c: i, glm interaction treatment, stimulus, group: p=0.038, d=0.464), again confirming previous data^18,19^. As Dnmts were also found to promote extinction learning^20^, we exposed the bees to the CS+ 6 times (Fig. 1a). We found that inhibiting Dnmts with RG108 led to less extinction across all 6 stimuli (Fig. 1c: ii, glm interaction treatment, stimulus, group: p=0.019, d=0.243). Taken together, our measurements of M17 responses in our preparation confirmed previously published data, and showed that the experimental treatments used here (in particular, staining with FURA on day two, and keeping the bees in the recording chamber for three days) did not affect the bees’ capacity to learn, and did not modify the effect of Dnmts on memory formation and extinction.

### Dnmts promote fast odor identity processing following olfactory reward conditioning

We recorded odor responses in PNs 2 days after training (Fig. 1a,b). Since Dnmts have been implicated with odor discrimination after learning, we first analyzed how similar the responses to two different odors were. Thus, we calculated the Euclidean distance (i.e. dissimilarity measure) between the odor response patterns to the CS+ and a new odor 2 days after olfactory reward conditioning (Fig. 2a). Background dissimilarity (noise) was in the range of 0.05. Upon odor stimulation, the dissimilarity increased to above 0.1, and decreased slowly thereafter. In the paired group, Dnmt inhibition led to less distinct odor patterns upon stimulus presentation (Fig. 2a). In the unpaired group, odor responses were more distinct in the Dnmt-inhibited group, with a slower return to baseline. The most prominent effect of Dnmt inhibition was in the initial odor response (Fig. 2b). With active Dnmts during memory formation the Euclidean distance increased in the paired group within the first 81-160 ms (Fig. 2c, glm interaction treatment, group: paired: p=0.018, d=1.408, unpaired: p=0.951). When averaged across the whole odor period, there was no significant difference between treatments (Fig. 2c, glm interaction treatment, group: paired: p=0.273; unpaired: p=0.076).

**Figure 2:**
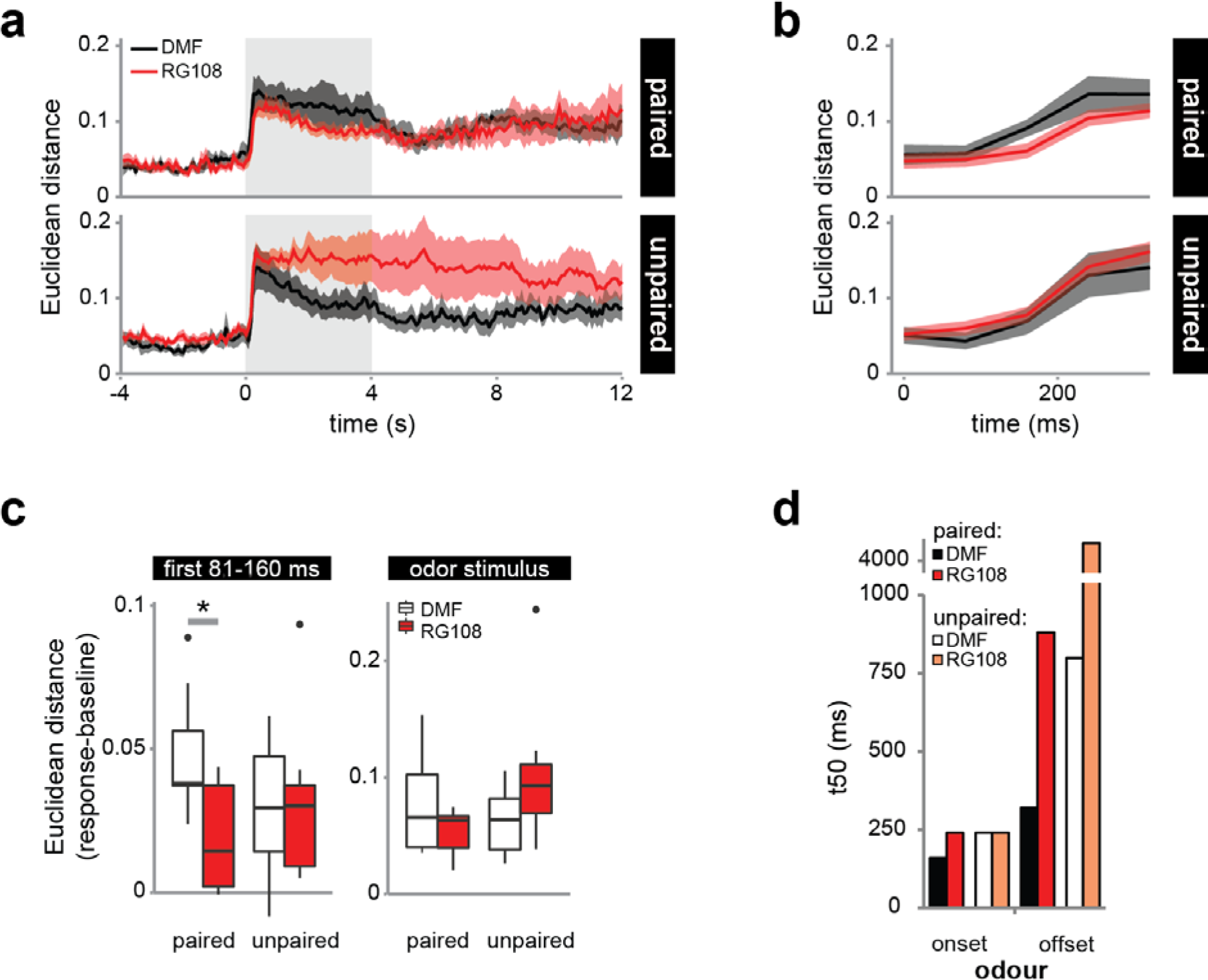
Dnmts accelerate fast odor response pattern separation. (a) Odor stimuli (shaded area) elicited significant responses in all groups (Euclidean distance as dissimilarity measure, mean +/− SEM). (b) Odor-specific patterns established at a slower pace when Dnmts were inhibited: 81-160 ms after odor onset dissimilarity was smaller in the paired/RG108 group. (c) The Euclidian distance was significantly different between treatments in the paired (glm interaction treatment, group: p=0.018, effect size (d)=1.408), but not the unpaired group (p=0.951). There was no difference when considering the entire odor stimulus (glm interaction treatment, group: paired p=0.273; unpaired p=0.076). (d) Training led to faster establishing (onset) and abolishing (offset) of distinct odor response patterns (paired vs. unpaired). Odor response pattern dynamics were also faster with active Dnmts (DMF) compared to inhibited Dnmts (RG108) in trained bees. Number of bees: paired: n(RG108)=10, n(DMF)=7; unpaired: n(RG108)=10, n(DMF)=6; * is p<0.05

How fast does the AL build its spatial odor representation? We calculated the time-point after odor onset when the majority of bees displayed distinct odor response patterns (t50-onset, see methods, Fig. 2d). When bees were trained to an odor, this time decreased from 240ms to 160ms. However, this effect was abolished when Dnmts were inhibited (Fig. 2d). The contribution of Dnmts was even larger for odor offset: pattern similarity in the majority of solvent treated paired bees returned to baseline levels after 320 ms, against 880 ms in RG108 treated paired bees. Again, RG108 treated paired bees were more similar to unpaired control bees (DMF unpaired: 800 ms; RG108 unpaired: 4280 ms). We conclude that Dnmts modify the antennal lobe neural networks during memory formation in a way that could allow for faster odor pattern discrimination.

### Dnmt activity during memory formation increases the number of glomeruli responding to a new odor

Odor learning can change the odor response strength in olfactory glomeruli depending on their activity during training^24^. Therefore, we quantified the percentage of activated glomeruli^24^. More glomeruli responded to a new odor after learning than to the CS+ (Fig. 3a). However, this effect was reversed when Dnmts were inhibited during memory formation: fewer glomeruli responded to the novel odor, as compared to control bees (Fig. 3a, glm interaction Treatment, Group, Stimulus: p=0.002, d=2.248). The number of odor-activated glomeruli in response to the new odor did not change when Dnmts were blocked in the unpaired group, where no odor learning took place (Fig. 3b, glm, p=0.495). This analysis shows that Dnmts are involved in recruiting intermediately active glomeruli into the responses to new odors after learning.

**Figure 3:**
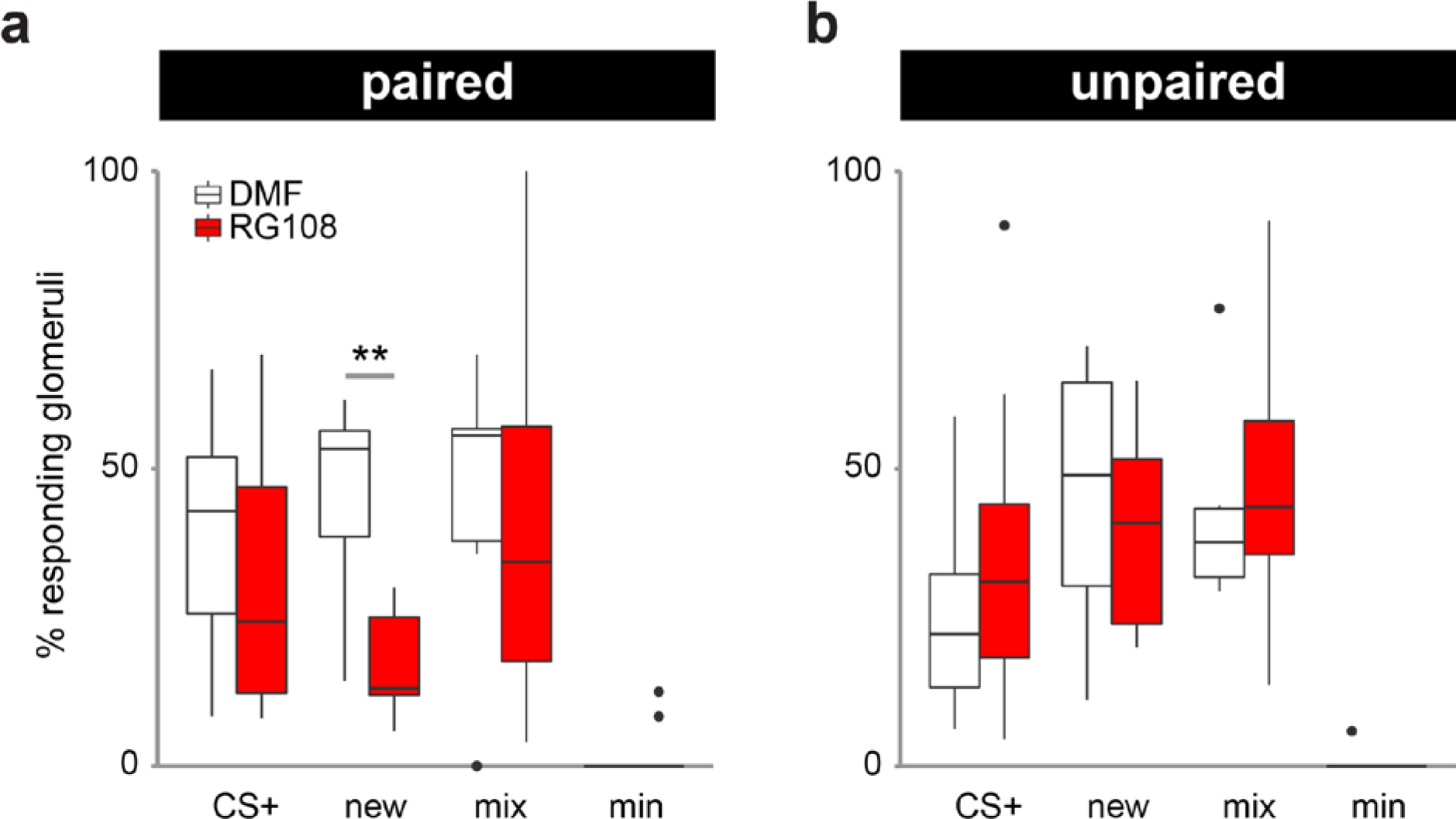
Dnmt activity during memory formation increases the number of glomeruli responding to a new odor. For each bee the % of glomeruli responding to each odor stimulus is plotted. Responding glomeruli were defined using the same criterion as described before^24^. (a) Less glomeruli responded to the new odor after RG108 treatment in the paired group (glm, interaction treatment, group, stimulus: p=0.002, effect size (d)=2.248), (b) but not in the unpaired group (p=0.495). Number of bees: paired: n(RG108)=10, n(DMF)=7; unpaired: n(RG108)=10, n(DMF)=6; ** is p<0.01

### Dnmt activity during memory formation leads to stronger response in new odor glomeruli

How are odor responses modified in the dominant glomeruli of each odor-response pattern? We focused on the two dominant glomeruli for response to the CS+ and the new odor, respectively (Fig. 4). The dominant CS+ glomeruli showed weaker responses to the new odor, and intermediate responses to the mixture of the two odors (Fig. 4a). Blocking Dnmts led to a slower return to baseline in the CS+ response in both paired and unpaired bees, and to the mixture in unpaired bees. Averaged over the entire odor response there was no significant effect of DNA methylation for paired bees, whereas Dnmt activity led to weaker responses to the CS+ in unpaired bees (Fig. 4b, glm interaction treatment, stimulus, group: p=0.042, d=0.775). Glomeruli most responsive to the new odor showed an increased response to this odor during the entire stimulus period when Dnmts had been active during memory formation (Fig. 4c). This difference was significant in the paired group, while there was no difference in the unpaired group (Fig. 4d, glm interaction treatment, stimulus, group: paired: p=0.024, d=0.938, unpaired: p=0.525).

**Figure 4:**
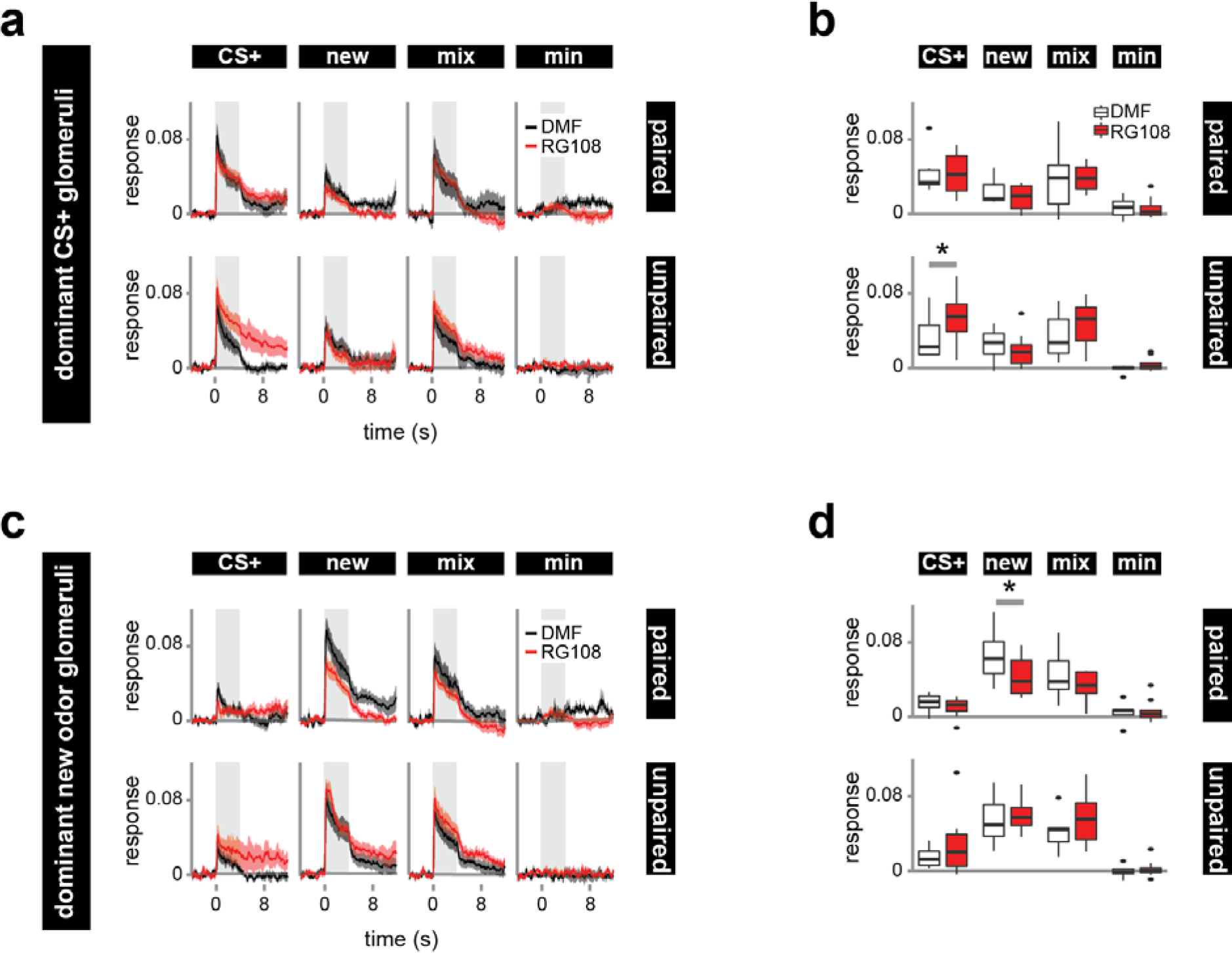
Dnmt activity during memory formation increases the response strength of glomeruli strongly responding to the new odor. Response strength of glomeruli was assessed by analyzing those two glomeruli responding most to the (a,b) CS+ and (c,d) the new odor respectively (dominant glomeruli). (a) For the dominant CS+ glomeruli, the average response over time and (b) pooled across the 4s odor stimuli is shown. The responses did not significantly differ between RG108 (red) and DMF (black) treated bees in the paired group. In the unpaired group, however, the response to the CS+ increased in RG108 treated bees (glm interaction treatment, group, stimulus: p=0.042, d=0.775). (c,d) The responses of dominant new odor glomeruli differed between treatments in the paired group. (d) The response was weaker in RG108 treated bees when stimulated with the new odor (glm interaction treatment, group, stimulus: p=0.024, effect size (d)=0.938). This, however, was not the case in unpaired bees (p=0.525). (a,c) The mean (+/−SEM) is shown. The shaded area indicates the odor stimulus. Number of bees: paired n(RG108)=10, n(DMF)=7; unpaired n(RG108)=10, n(DMF)=6; * is p<0.05

### Dnmt activity during memory formation does not affect odor responses in the AL during extinction learning

Extinction is a form of learning that occurs when a learned odor is presented repeatedly without reinforcement. Extinction can be influenced by Dnmts^20^. We exposed bees to the CS+ six times following the odor test (Fig. 1a). We calculated how the representation of the CS+ changed during these six presentations by calculating the Euclidean distance to the first presentation: a value of 0 indicates that the odor response remained stable. We found that odor responses changed slightly with accumulating extinction trials in the paired group (Fig. 5a, glm interaction treatment, stimulus, group against 0: 5^th^ trial RG108: p=0.012, d=0.777, 6^th^ RG108: p=0.042, d=0.714, 6^th^ DMF: p=0.005, d=0.754), but without a difference between treatments. Response variability was higher in the unpaired group, but only when Dnmts where inhibited; here the odor responses differed significantly from the first presentation in all extinction trials (Fig. 5a, glm interaction treatment, stimulus, group against 0: 2^nd^ trial: p=0.009, d=0.838, 3^rd^: p=0.035, d=0.731, 4^th^: p<0.001, d=0.801, 5^th^: p<0.001, d=0.705, 6^th^: p=0.007, d=0.639), with the odor patterns being more distinct in RG108 compared to DMF treated bees in two trials (Fig. 5a, glm interaction treatment, stimulus, group: 4^th^ trial: p=0.039, d=0.984, 5^th^: p=0.034, d=0.896).

**Figure 5:**
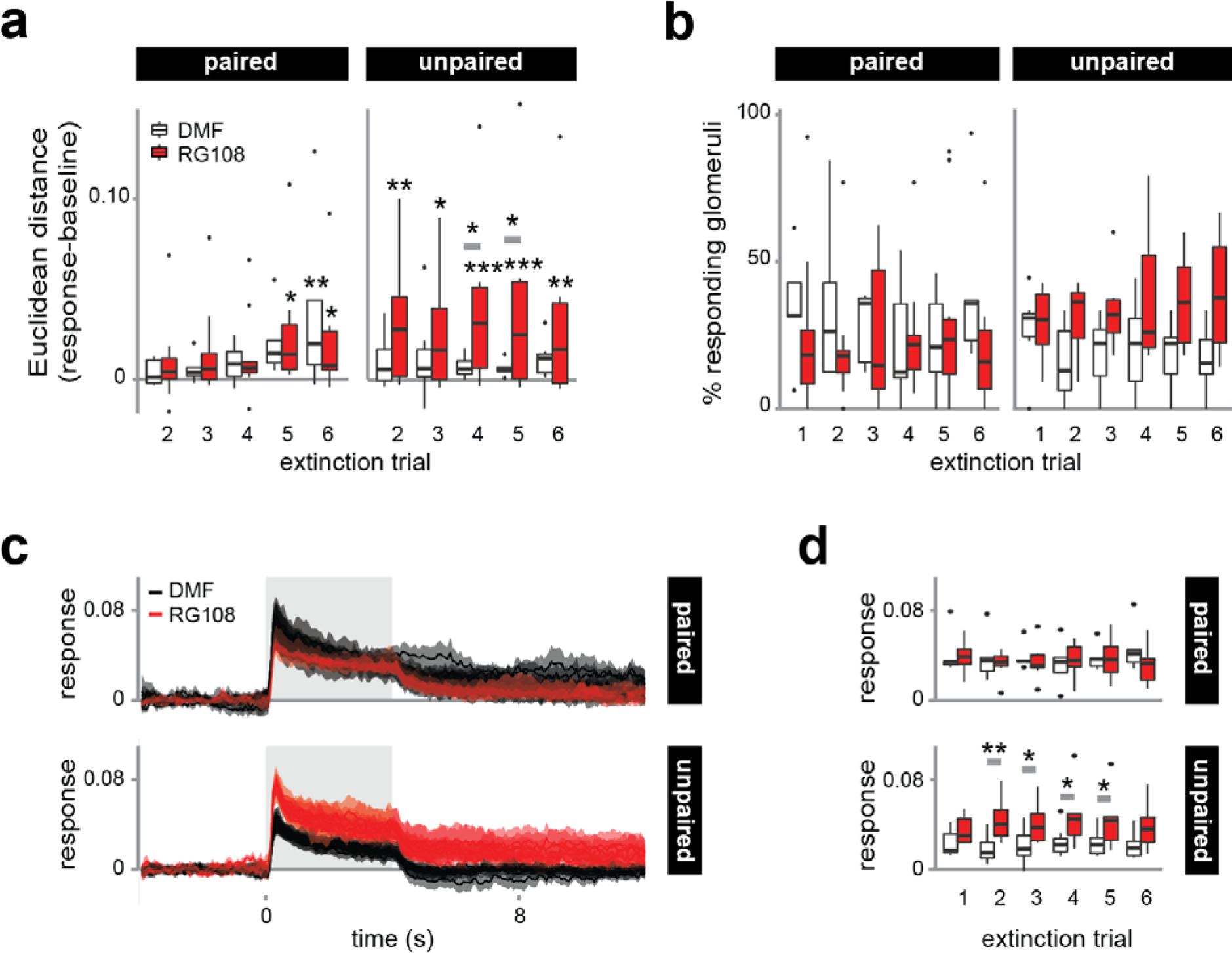
Dnmts do not affect odor responses during extinction learning in the AL. Bees were exposed to the CS+ six times, which causes extinction learning and a reduced PER response. (a) the Euclidean distance (dissimilarity measure) in relation to the first extinction trial differed between RG108 (red) and solvent (white) treated bees only in the unpaired group (glm interaction treatment, group, stimulus: 4^th^ trial: p=0.039, d=0.984, 5^th^: p=0.034, d=0.896). (b) The number of active glomeruli did not change for any treatment or group. (c) The averaged response (+/−SEM) of the dominant CS+ glomeruli is shown over time for all six trials. In paired bees the response was similar in RG108 (red) and DMF treated bees (black). In unpaired bees, however, the response during and after the odor stimulus is increased after Dnmt inhibition. (d) Pooled across the entire 4s odor stimulus, response strength increased following Dnmt inhibition for four out of six extinction trials in the unpaired group (glm interaction treatment, group, stimulus: 2^nd^ trial: p=0.007, d=1.573; 3^rd^: p=0.038, d=1. 149; 4^th^: p=0.016, d=1.058; 5^th^: p=0.030, d=1.025). Number of bees: paired n(RG108)=10, n(DMF)=5; unpaired n(RG108)=6, n(DMF)=7; * is p<0.05; ** is p<0.01; *** is p<0.001

The number of responding glomeruli did not change for either treatment or group (Fig. 5b). However, the two dominant glomeruli increased their response strength to the repeated stimulus in the unpaired/RG108 group (Fig. 5c). This effect was significant in the 2^nd^-5^th^ extinction trial (Fig. 5d; glm interaction treatment, stimulus, group: 2^nd^ trial: p=0.007, d=1.573; 3^rd^: p=0.038, d=1.149; 4^th^: p=0.016, d=1.058; 5^th^: p=0.030, d=1.025). Together, our data indicate that glomeruli responses remained largely stable over repeated odor stimulations. At least in the unpaired group (no learning, but pre-exposure) this stability necessitates a Dnmt-dependent mechanism.

## Discussion

Here we investigated whether and how Dnmts mediate plasticity in the honeybee AL after olfactory reward conditioning and during extinction. Using Ca^2+^-imaging of odor evoked PN activity in the AL we show that Dnmt-mediated DNA methylation during memory formation specifically promotes the number and response strength of glomeruli responding to a new odor 2 days after training. Additionally, the dynamics of odor pattern separation between the CS+ and a new odor changed depending on Dnmt activity during memory formation. Furthermore, AL responses during extinction learning were not affected by Dnmt activity. After stimulation (i.e. unpaired training) alone, however, Dnmt activity during memory formation promoted a stable response of glomeruli with repeated stimulations of the pre-exposed odor.

### Dnmts may promote stimulus-specific memory formation by facilitating fast odor pattern separation

The findings described here can be directly connected to what we know about Dnmts function in memory formation from behavioural studies. Dnmts promote stimulus-specific LTM formation after olfactory reward conditioning^18,19^. We could show here that Dnmts also promote fast odor response pattern separation between the CS+ and a new odor. Odor discrimination in the AL is fast and maximum pattern separation is reached around 150 ms after odor onset in PNs^7,8^. Bees respond behaviourally to trained odors within 430-470 ms^9,10^. Furthermore, bees can successfully discriminate odors, even if they smell them for just 200 ms^9^. This suggests that bees use information about odor pattern similarity which is generated during the first few hundred milliseconds, in order to decide whether to respond to an odor or not. An associative change in the temporal dynamics of odor pattern separation-mediated by Dnmts-would have a strong impact on generalization between odors and thus stimulus-specific memory.

### Dnmts might affect memory-related plasticity in local inhibitory neurons of the AL

Interestingly, Dnmt activity during memory formation did not globally affect response strength; it rather specifically increased the number and strength of glomeruli not strongly active during the training. Our results allow speculation that Dnmts might regulate the strength of inhibitory connections from CS+ glomeruli to those weakly active or inactive during training (Fig. 6). The majority of inhibitory LNs in the AL are heterogeneous, branching strongly in one glomerulus and weakly in few others^25,26^. Indeed, the glomeruli most active in response to the two odors used here have inhibitory connections onto each other^2,27^. Additionally, it has been suggested earlier that heterogeneous LNs are plastic following olfactory reward learning^3,28^ and that they play a crucial role in odor discrimination^4,5^. Alternatively, or additionally, synaptic plasticity might occur in a glomerular subpopulation of output synapses in homogeneous neurons, yielding a spatially complex functional pattern.

**Figure 6:**
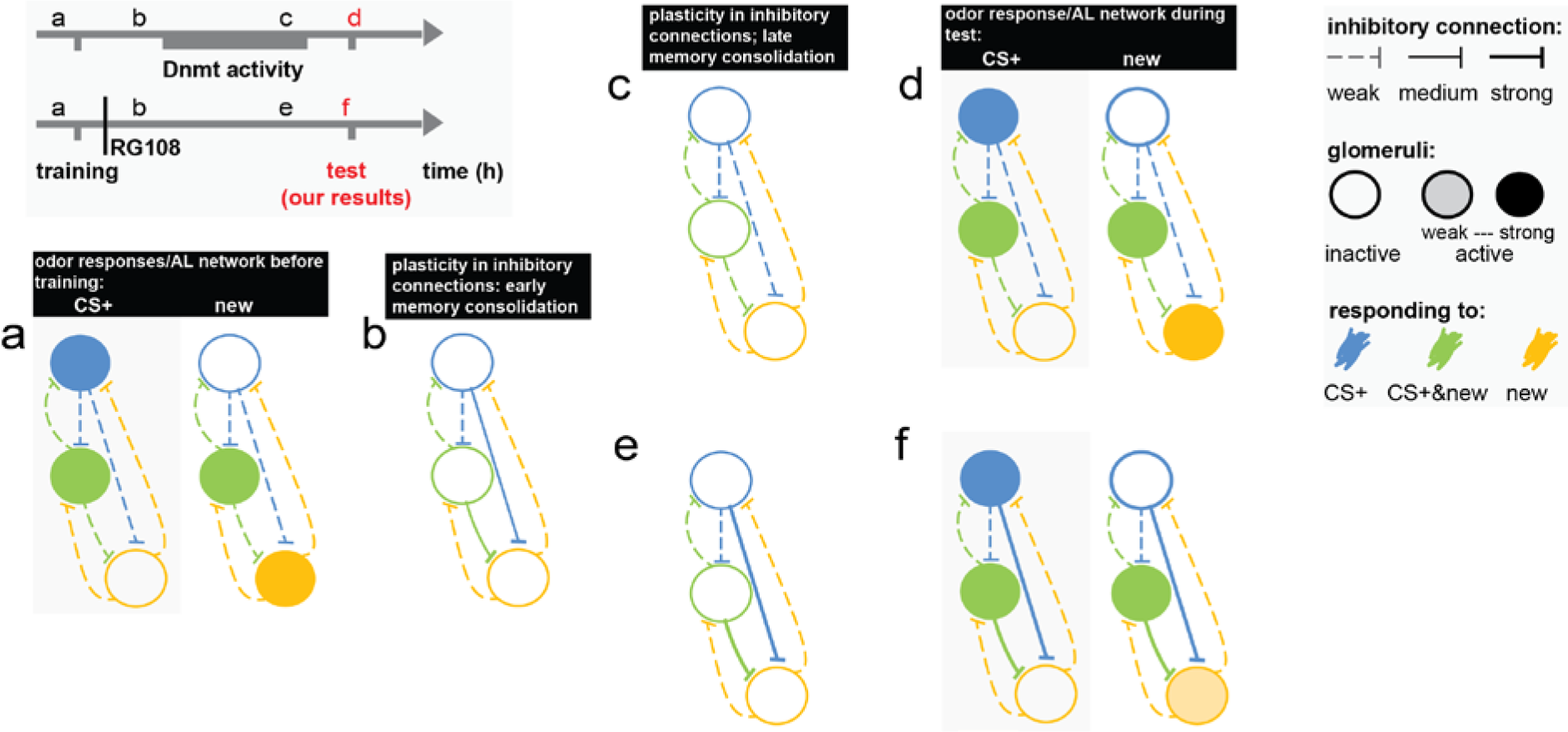
Dnmt mediated regulation of the AL LN network could explain the effects observed here. Our results show that Dnmt activity during memory formation affects the response of glomeruli, which respond strongly to a novel odor but weakly or not to the CS+. Plasticity in the response of these glomeruli may result from changes in the inhibitory local interneuron (LN) network within the AL. We suggest the following model: (a) assume a simplified AL, with one glomerulus (blue) responding only to the CS+, one (yellow) not to the CS+, but to another odor, and one (green) responding to both. A (weak) network of inhibitory LNs connects all glomeruli. (b) When an odor (CS+) is paired with a reward (US), the representation of that odor is initially strengthened by increasing the inhibitory connection to nonactivated glomeruli (from blue and green onto yellow). (c) These inhibitory connections are reduced again by a homeostatic mechanism mediated by DNA methylation, avoiding that the AL network saturates over many learning events. (d) As a result, during the odor test the overall network inhibition would be recovered. (e) In the absence of Dnmt activity inhibitory connections remain strong or further increase in strength, (f) therefore reducing the response of glomeruli to a novel odor during the test 2 days after training. The model assumes that learning induces the growth of new (inhibitory) synapses, and DNA methylation mediates the termination of this growth by downregulating memory-associated genes such as *Actin* and/or homeostatic plasticity mechanisms such as synaptic scaling^21,22^

Dnmt activity is associated with decreased expression of memory-associated synaptic genes (e.g. *actin* and *neurexin I*)^19^, after initial expression waves during the first hours after training^29^. This process might be important for restricting synaptic plasticity in the LN network after olfactory reward training (Fig. 6), creating a temporal window for learning induced synaptic changes, followed by a temporal window for homeostatic regulation.

If Dnmts predominantly regulate plasticity in the AL LN network, then the contribution of Dnmt activity to olfactory memory specificity should depend on the degree of inhibitory connections between the glomeruli responding to the CS+ and a new odor. Therefore an important next step is to test the relationship between associative plasticity in the inhibitory local AL network and Dnmt activity, ideally by recording directly from LNs.

### Dnmts might serve distinct regulatory functions following learning and odor exposure

Here we investigated the role of Dnmts in both animals which formed memories (i.e. paired) and those which were stimulated with odor and sugar repeatedly, but did not form memories (i.e. unpaired) during training. The differences we found between these two groups highlight two interesting aspects of how Dnmts might regulate transcription-dependent plasticity in the AL: (1) part of the regulation is memory-dependent. This is supported by evidence that some memory-associated genes show learning-dependent changes in their methylation pattern^19^. (2) Dnmt-dependent plasticity after learning and stimulation had different characteristics, as in one case the immediate response to a new odor changed, and in the other the repeated response to the pre-exposed odor. This suggests that DNA methylation may have two distinct roles in this context: first; to restrict gene expression levels during memory formation^19^ and second; to regulate re-expression of genes^30^. This may explain why different genes are targeted by Dnmts under different circumstances: in some genes DNA methylation changes occur exclusively in response to learning, and in others in response to both learning and stimulation or to stimulation only^19^.

### Dnmts might contribute to homeostatic plasticity mechanisms acting on the level of whole cells and neural networks hours and days after training

Different types of plasticity can occur following neural activity, including immediate Hebbian and protracted homeostatic plasticity^31^. Homoeostatic plasticity globally counteracts activity induced local changes in order to normalize overall activity levels and prevent extrema^31–33^. Homoeostatic plasticity is induced by and utilizes mechanisms (e.g. intracellular Ca^2+^ levels) which are partly the same as those responsible for long-term potentiation (LTP, i.e. the cellular equivalent of LTM)^31^. The important distinction, however, lies in the time-scale they are acting on, as homeostatic plasticity operates within hours and days, instead of seconds^31,33^. Furthermore, in contrast to local synapse-specific changes, homeostatic plasticity acts globally on the whole cell or neural network. Neural network models suggest that homoeostatic plasticity is important for counteracting accelerating activity by preventing positive feedback loops and for remaining plastic^34,35^. Recent evidence suggests that DNA methylation levels can control synaptic scaling, a mechanism of homeostatic plasticity, *in vitro* in mammals^21,22^. In bees, DNA methylation might also regulate homeostatic plasticity: *Dnmts* are upreglated on a time-scale corresponding to that of homeostatic rather than Hebbian plasticity^19^. Furthermore, Dnmts are likely involved in the downregulation of a subset of memory-associated genes during memory formation^19^. We earlier proposed that Dnmts-in particular Dnmt3-may act by normalizing the expression patterns of target genes following an initial upregulation after training^19^. At the molecular level, such a process could contribute to homoeostatic plasticity aiming at reducing overall cell activity, excitability and synaptic growth back to baseline levels by downregulating transcription. Our results suggest that plasticity in inhibitory LNs might be mediated by Dnmts (Fig. 6). Inhibitory neurons perform a crucial function for network homeostasis, as they regulate overall activity in a network by adjusting inhibition^33^. Considering DNA methylation and homeostatic plasticity may be directly connected, these observations provide a credible starting point to address the role of epigenetic transcriptional regulators in governing the dynamics of neural networks during memory formation.

## Online Methods

### Olfactory training and treatment

Honey bees *(Apis mellifera)* were trained using appetitive olfactory classical conditioning as described before^18,19^. In short, bees received six trials of odor (conditioned stimulus, CS) and sugar (unconditioned stimulus, US) pairings. In one group of bees the CS and US overlapped for 2 s (i.e. paired group), which causes stable long-term memory formation. Another group of bees received the CS and US with a 5 minute gap between the stimuli (i.e. unpaired group), which does not cause long-term memory formation or conditioned inhibition^36^. In the paired group bees responded to the CS+ on average in 4.5 out of 6 training trials and in the unpaired group in 0.15 out of 6 (Tab. 1). Both groups were trained in parallel, to avoid the influence of seasonal and day-to-day variability. In both cases the odor stimulus lasted 4 s and the sugar stimulus (1 M sugar water) 3 s. Bees were either trained with 1-hexanol or 1-nonanol (10^2^ in mineraloil, all Sigma-Aldrich, St. Louis, USA). 100 μl of the diluted odor was applied to a cellulose stripe (Kettenbach GmbH KG, Eschenburg, Germany) located in a 3 ml syringe (Henke-Sass, Wolf GmbH, Tuttlingen, Germany). 2 hours after training 1 μl of the Dnmt inhibitor RG108 (Sigma-Aldrich, St. Louis, USA, 2 mM in DMF) or the solvent DMF was applied topically on the thorax. This treatment method successfully reduces DNA methylation in the brain^19^. From two hours after training, bees were repeatedly fed to saturation with 1M sugar water until the night before Ca^2+^-imaging or M17 recordings.

### Projection neuron staining and imaging

24 hours after training, the bees’ lateral and medial antenno-protocerebral tracts (land m-APT) were stained with Fura-2 dextran (Invitrogen, ThermoFisher Scientific, Waltham, USA) - a Ca^2+^-sensitive dye - by inserting a dye crystal with a glass electrode between the MB calyces. Staining and preparation for imaging was done as described before^24^, with minor alterations. The brain was covered with bee saline solution (NaCl 130mM, KCl 6mM, MgCl_2_ 4mM, CaCl_2_ 5mM, Sucrose 160 mM D-Glucose 25 mM, HEPES 10 mM, pH 6.7). The bees were kept overnight in a humid plastic container. Imaging of bees started 2 days after training. As 16-48 bees were trained each day the actual time between training and imaging for each individual bee differed. On average bees were imaged 52 hours after training (for more information see: Tab. 1). A total of 40 bees were imaged and analysed (DMF paired: 9; RG108 paired: 13; DMF unpaired: 8; RG108 unpaired: 10). Bees were imaged as described before^24^ with a spatial sampling rate of 172 × 130 pixel, using a 20x dip objective (NA=0.95), and a Till-Imago CCD camera. Each recording lasted 16 s (200 frames) with the odor stimulus starting 4 s into the recording and lasting 4 s. Each frame was recorded with 340 and 380 nm excitation light at a rate of 12.5 Hz, thus one double frame lasted 80 ms. Odors were delivered during the measurement as described before^24^. Bees received an odor test first, consisting of the CS+, a new odor and mineraloil in randomized order followed by the binary mixture of CS+ and new odor (Fig. 1a). The odor test was followed by an extinction paradigm, consisting of a presentation of the CS+ 6 times and one presentation of mineral oil (solvent stimulus) at the end of the measurement as a contamination control. Stimuli were separated by 1 minute.

**Table 1.** Overview over acquisition scores, measurement time, AL and glomeruli analysed. For each group the mean (+/−SEM) across all bees used in the imaging experiment is shown for the accumulated CS+ response during 6 training trials, the time between training and imaging, the AL imaged and the number of glomeruli analysed.

|  | paired |  | unpaired |  |
| --- | --- | --- | --- | --- |
|  | DMF | RG108 | DMF | RG108 |
| **Training PER (max. 6)** | 4.7 (+/-0.4) | 4.3 (+/-0.5) | 0.1 (+/-0.1) | 0.2 (+/-0.2) |
| **Time after training (h)** | 51.7 (+/-0.5) | 51.7 (+/-0.8) | 52.4 (+/-1.0) | 51.3 (+/-0.8) |
| **AL (1=left; 0=right)** | 0.4 | 0.6 | 0.9 | 0.5 |
| **Number of Glomeruli** | 14.2 (+/-1.0) | 15 (+/-1.5) | 15.5 (+/-2.5) | 15 (+/-2.0) |

### M17 recordings

For M17 recordings, bees were stained as described above. 48 hours after training the M17 response was recorded. M17 activity correlates with the PER^23^ and can therefore be used to assess memory retention. One 0.2 mm insulated silver wire was inserted between the bees’ compound eye and lateral ocellus into the muscle, and a second one in the opposite eye as a reference (Fig. 1b). The signal was detected by a custom built digital oscilloscope with a resolution of 0.0625 ms and was connected to the electrodes via an amplifier. Baseline spike frequency was measured for 5 s before every odor stimulus. Immediately afterwards spike activity during the 4 s odor stimulus was recorded. The odor stimuli were the same as described above and shown in Fig. 1a. After the measurement, the bee’s PER was elicited by stimulating the antennae with 1 M sugar solution. All bees in which the M17 did not show activity in response to sugar were excluded. In sum, 73 bees were measured and analysed (DMF paired: 26; RG108 paired: 19; DMF unpaired: 13; RG108 unpaired: 15).

### Data analysis

All data analysis except the pre-processing of imaging data was done in R^37^. All scripts were custom written. M17 data was analysed by extracting the number of spikes during the 5 s baseline and during the 4 s odor stimulus period. The M17 response frequency was calculated for each odor stimulus and was normalized with the corresponding baseline frequency.

Imaging data were pre-processed with the ImageBee plugin for KNIME^38^ Movement correction was performed for each bee first between images (i.e. frames) and then between videos (i.e. stimuli). Signals were calculated as the ratio of fluorescence at 340 and 380 nm: *F*_340/380_ = ^*F*^_340_/*F*_380_. The *F_340/380_* was then normalized to baseline levels by subtracting the average *F_340/380_* of the first 40 frames (i.e. before odor onset). For glomeruli detection, videos were processed as follows: A Z-score normalization was performed, images were smoothed with a Gaussian filter, a principal component analysis was run and a convex cone algorithm was used as described elsewhere^38^. The map of glomeruli obtained by this procedure was than overlaid with the *F_340/380_* calculations. The response of each glomerulus over time was calculated by averaging all pixels in the identified area. On average, 15 glomeruli could be analysed per bee (for more detail: Tab. 1). Bees which showed strong movement during one of the stimuli were excluded from the equivalent part of the analysis (i.e. test or extinction).

We calculated the Euclidean distance from the glomerular responses^39^ for each individual bee. We determined t50 as the time point after odor onset (and offset) were the Euclidean distance in more than 50% of bees (less than 50% of bees) exceeded 3x the standard deviation (SD) of the baseline period.

We determined the glomeruli responding to each stimulus as described before^24^. All glomeruli exceeding 3x SD of the period before odor onset were counted as responsive. We determined the two most active glomeruli during the peak response for the CS+, new odor and first extinction trial for each individual bee. We pooled the response of those two glomeruli and calculated the mean and SEM across bees. We assessed the two strongest instead of all responding glomeruli as this method avoids introducing a bias caused by the reduced number of active glomeruli in the new odor after RG108 treatment (Fig. 3). Additionally, as each individual bee was trained with either 1-hexanol or 1-nonanol, the identity of the CS+ and new odor was different across bees. These two odors differ in which and how many glomeruli are activated^2,40,41^.

We used generalized linear models for statistical significance testing. The effect size (Cohen’s D) was calculated for all effects reaching the 0.05 significance level. As a guideline effects with sizes below 0.2 were defined as negligible, between 0.2 - 0.5 as small, between 0.5 - 0.8 as medium and above 0.8 as large^42^. The effect size can be used as an estimate of the real difference between the tested groups.

## Acknowledgements

We thank Paul Szyszka, Stefanie Neupert and Nicholas Kirkerud for their feedback and technical support. The study was funded by grants from the Australian Research Council and National Health & Medical Research Council of Australia (ARC DP120104117, ARC DP120102301, NHMRC APP1008125 to CC and JR), and by the Deutsche Forschungsgemeinschaft DFG (SPP 1392 to CGG). CC was also supported by an Australian Research Council Future Fellowship (FT110100292). SDB was funded by an International Postgraduate Research Scholarship, University of Queensland Centennial Scholarship and graduate school international travel award.

## Author’s contributions

SDB, CC, JR and CGG conceived the study. SDB and CGG designed the experiments. SDB performed the experiments. SDB analysed the experiments. SDB wrote the paper. CC and CGG edited the paper.

